# Intricate asymbiotic hyphal interactions of *Rhizophagus irregularis* revealed with single-plane observations in a new microfluidic device

**DOI:** 10.64898/2026.02.17.706285

**Authors:** Felix Richter, Erica McGale, Maryline Calonne-Salmon, Kate Collins, Stéphane Declerck, Ian R. Sanders, Claire E. Stanley

## Abstract

Arbuscular mycorrhizal fungi (AMF) are essential symbionts to most land plants. There is great interest in using AMF inoculation treatments to enhance restoration or agricultural efforts. However, little is known about AMF asymbiotic interactions that could shape soil ecosystems and inoculation outcomes. Studying these early traits is challenging given the obligate nature of AMF. Microfluidic devices enable high-resolution, single-plane, time-resolved observation of AMF hyphal traits in controlled, plant-less environments. The *AMF-AnastomosisChip*, introduced in this work, additionally separates spores and hyphae into lanes for detailed observations prior to entering a shared interaction zone. Using this device with *Rhizophagus irregularis*, we phenotyped key hyphal traits from germination to anastomosis. We compared the effect of a fatty acid (FA) treatment to other environmental factors in shaping AMF growth and fusion (anastomosis). Among our results, we confirm known effects of the FA treatment in increasing hyphal branching and longevity but interestingly show that multiple hyphae in a lane can negate FA effects. We also newly reveal that anastomosis occurs for 100% of interactions among hyphae from different lanes, but only for 50% of in-lane interactions, potentially linked to spore origin. Our findings validate the *AMF-AnastomosisChip* as a versatile platform for increasing discovery in trait-based microbial ecology, including for asymbiotic AMF interactions with environmental factors. This work sets the foundation for future studies with the *AMF-AnastomosisChip* on the effects of nutrient content, plant-derived molecules, or microbial community members on the AMF traits characterized in this study.

## INTRODUCTION

The symbiosis of arbuscular mycorrhizal fungi (AMF) with over 70% of all land plants lays the foundation for all soil-based ecosystems on the planet [1–3]. AMF interactions with plants and soil are important for ecosystem stability [4,5], plant evolution [6], and agriculture [7]. Due to their key role in natural and agricultural ecosystems, AMF have been the subject of many investigations in the last decades. Nevertheless, much remains unknown about these fungi, and especially non-plant drivers of AMF growth in the asymbiotic and symbiotic phases.

AMF observational research, which is necessary to dissect the influences on their traits, is challenging. As AMF are obligate symbionts, it is difficult to maintain plant-less AMF cultures long enough to observe the intricacies of asymbiotic growth, though some researchers have succeeded under certain conditions [8–10]. To date, the association of AMF with root organs is a well-established method for their *in vitro* cultivation until spore production and allows detailed observation of hyphal growth throughout their lifespan, though this is purely during the symbiotic phase [11–14]. Hyphae can be observed alone in media in bi-compartmented plates: the plant-AMF association is housed on one side of the plate and hyphae then pass over the central divider (*e.g.*, in root-organ culture) to continue growth in media without plant root exudates, nor sucrose [15,16]. This set-up has led to several inter- and intra-specific hyphal trait screenings [17–19]. Still, visualization of symbiotic-phase hyphal interactions, let alone those from the asymbiotic phases, remain difficult due to the multi-plane nature of solid media.

Apart from the challenge of combining hyphal growth and interaction data across planes, observing hyphal networks in solid media is fundamentally different than in soil. Liquid media may more closely allow for natural hyphal growth and exploration than solid media, but this is not entirely known. With the axenic cellophane sandwich method, which confines spores and their new hyphae in the z direction and uses liquid only to moisten the cellophane paper, microscopy observations and their interpretation can be improved significantly [8,20]. This has led to revelatory work where hyphae were confronted with rudimentary physical obstacles [21] or with different AMF strains [8,22]; importantly, both studies provided asymbiotic phase insights. Open questions remain on when (*e.g.*, at what distance) hyphae begin to change their growth in relation to encounters with abiotic and biotic environmental factors. Whether hyphae instead happen upon obstacles or other hyphae stochastically would be important for understanding the spatial scale at which AMF function.

Recently, a microfluidics-based approach to investigate asymbiotic hyphae was established. This provided increased insight into asymbiotic hyphal space searching mechanisms, the reversibility of cytoplasmic retraction, and hyphal growth prior to anastomosis [10]. By utilizing microfluidic technology, the microchannel height can be tuned precisely to the AMF studied and exploited to separate its different tissues; specific locations in the *AMF-SporeChip* were suited to spore diameters to trap them before germination (i.e., 100 µm microchannel height for *R. irregularis* spores), leaving only hyphae to grow into the investigation zone (*i.e.,* 10 µm microchannel height) for observation due to efficient z-confinement. Together with the near perfect transparency of the device materials (glass and the elastomeric polymer, poly(dimethylsiloxane) (PDMS)), this significantly enhances the quality of trait observation. Moreover, the microenvironment can be adapted to provide chemical niches or gradients to test hyphal habitat exploration and modification [23]. Even though the field of microfluidics is still relatively young, these advantages have already been applied to multiple fungal studies (reviewed in [24]).

As part of Richter and colleagues’ observations with the previous *AMF-SporeChip* [10], tip-to-side and tip-to-tip contact between hyphae from different individuals of the same strain versus from the same individuum could be separated. Potential pre-anastomosis signalling was observed in stop-and-go growth as hyphae approached other hyphae. Still, it was hard to exclude the influence of other local hyphae in each approach, with observed events being coincidental rather than controlled. To our knowledge, no studies, even using microfluidic technologies, have tracked asymbiotic, physically separated hyphae up to the point at which they have the chance to anastomose, in order to fully dissect their interaction mechanisms. Being able to screen these traits comprehensively could add information to databases such as TraitAM that can help answer biological, ecological, evolutionary and application questions related to spore, and more, traits of AMF [25].

We developed a new microfluidic device, tailored to study influential factors on AMF spore germinations and hyphal growth (direction, speed, branching) while isolated in standardized lanes. The device serves a second, equally important purpose: hyphae in the lanes are guided towards the interaction space, where directional changes at lane ends could be linked to properties of the interaction zone, *e.g.*, numbers of hyphae entering of the same or another isolate source. Altogether, this device allows dissection of hyphal growth traits leading to anastomosis and is, thus herein, introduced as the *AMF-AnastomosisChip*. It additionally allows the introduction of an abiotic or biotic treatment to characterize their influences on hyphal traits. In this study, the fatty acids (FAs) myristate and palmitate were introduced into our microfluidic chip set-up, since these compounds were both shown to enable and enhance AMF growth in asymbiotic growth conditions [9]. When considering hyphal interactions among different individuals or fungal isolates, introducing a FA treatment can putatively induce more competitive growth, given AMF need FAs produced by plants to survive and reproduce [26]. How FA-related changes in AMF hyphal growth and interactions may manifest can be tracked precisely in the *AMF-AnastomosisChip*. Additionally, these FAs were reported to lengthen the activity (defined throughout this manuscript as duration of growth) of asymbiotic AMF [9], allowing longer observations of hyphal traits in our devices.

Here, we studied the link between AMF growth and anastomosis when nearby spore and hyphal densities, previous traits of each hypha, and the presence of FAs is varied. We used the model AMF species *Rhizophagus irregularis* and introduced spores of the same isolate into physically segregated pockets, on opposite sides of the device, in the *AMF-AnastomosisChip*. We made detailed observations of AMF asymbiotic growth and interactions in a single plane. We used a variance decomposition analysis (repeatability) on spore germination rates, hyphal growth rates, hyphal exploration (lane crossing, branching), and anastomosis, to establish which experimental variables exhibited a significant influence on these traits. We show the importance of dissecting hyphal movement and interactions with physical separation, as in the *AMF-AnastomosisChip*, and reveal the use of this microfluidic device to further explore AMF ecological interactions, especially without the confounding influence of the plant host.

## MATERIALS AND METHODS

### Chicory root and AMF culture

The AMF strain *R. irregularis* MUCL 43194 (DAOM197198), was used for the experiment [27]. The fungus was maintained on root organ cultures of *Ri* T-DNA transformed roots of chicory (*Cichorium intybus* L., [12,28]). The original inoculum, *i.e.*, a plug with both the fungus and roots, was provided by the Glomeromycota *in vitro* Collection (GINCO, Belgium). This plug was used to make local cultures on Modified Strullu Romand (MSR) medium (pH 5.5) either with added sucrose (MSR+) or without (MSR-), solidified with 3 g/L phytagel. New chicory roots were grown on MSR+ medium in mono-compartmented Petri dishes (Ø = 90 mm) and subcultures were maintained by re-plating pieces every 1-2 months. Fresh and vital pieces of root were then used to inoculate bi-compartmented plates, in the growth format described by Rosikiewicz et al. [15] as the “standard culture system” for propagation and harvest of sterile, *in vitro* spores.

### Microfluidic device design and fabrication

The device design features two inlets and one outlet (Ø = 3 mm), with a footprint in the shape of a “T”, measuring 11.7 mm by 7.4 mm (Fig. 1A). The inlets are followed by a 1540 µm wide channel, which then opens slightly into an array of 9 equally spaced, rounded pockets on each side of the device (18 in total). These pockets are 170 µm at their widest point and connect into narrow lanes of 90 µm width and 400 µm length, which lead diagonally down into the central region, termed the hyphal interaction zone. This interaction zone, as well as the outlet channel below, are 500 µm wide. The device features two different microchannel heights (Fig. 1B). While the lanes and the hyphal interaction zone possess a channel height of 10 µm, the pockets and inlet/outlet channels are 100 µm high; this height difference facilitates trapping of the AMF spores [10].

**Figure 1.**
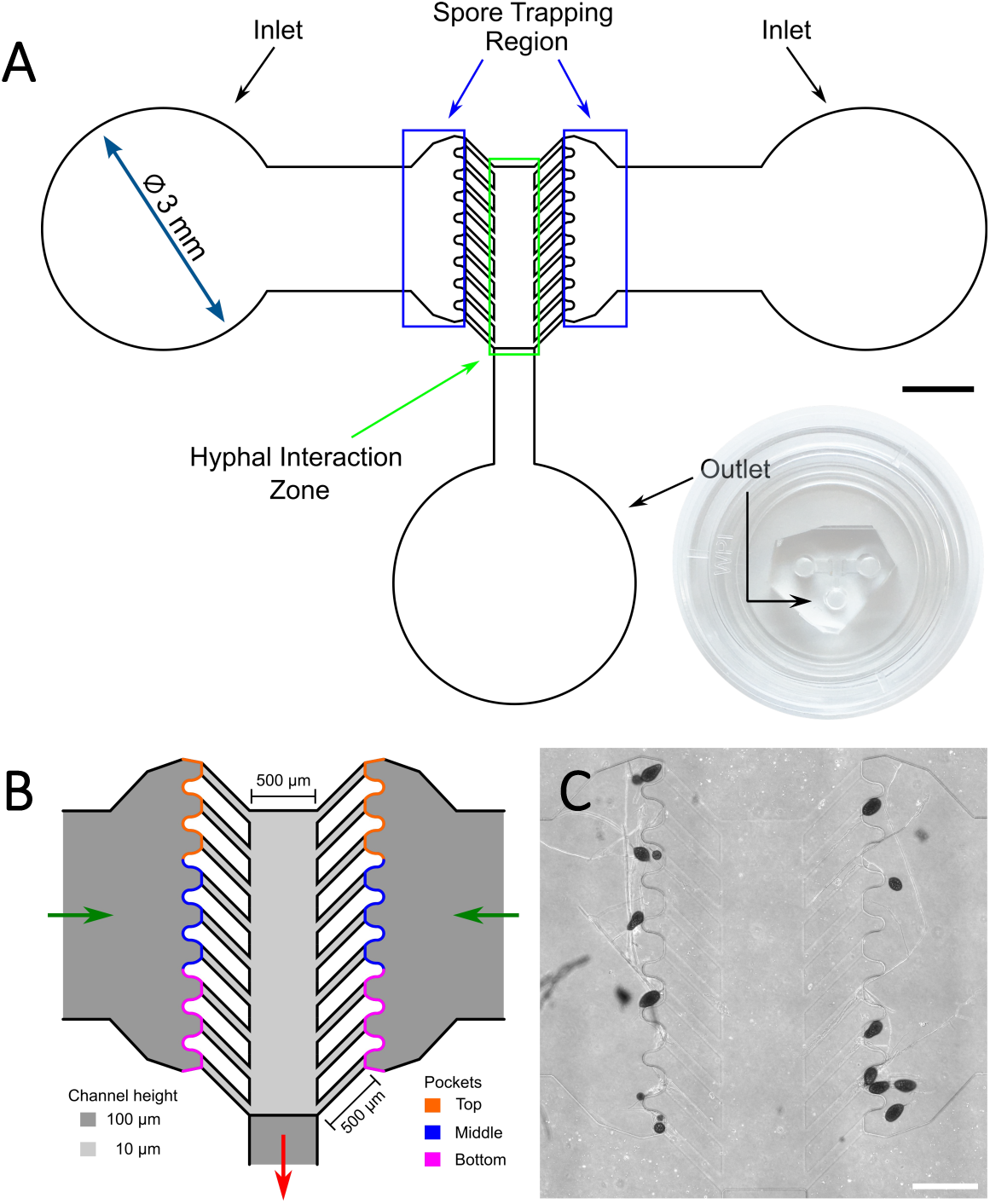
Design of the *AMF-AnastomosisChip*. **(A)** Schematic of the microfluidic device showing the dimensions and structure, highlighting the in- and outlets as well as the spore trapping region (blue boxes) and the hyphal interaction zone (green box). Inset: photo of the device. The scale bar only refers to the schematic. **(B)** Central region of the device. Dark grey indicates the in- and outlet channels, ending in the pockets, with heights of 100 µm; light grey indicates the lanes and interaction zone with a 10 µm height. Green arrows point out the direction of travel of the loaded, flushed spores from the inlets, while red arrows highlight the direction that excess media flows towards the outlet. To record loading data, the pockets are distinguished by their position in relation to the outlet: the three furthest from the outlet are considered the “Top” pockets (orange), then the “Middle” ones (blue), and then closest to the outlet are the “Bottom” pockets (pink). **(C)** Phase contrast image showing the distribution of *Rhizophagus irregularis* MUCL 43194 spores in the device. Scale bars: (A) 1 mm; (C) 400 µm.

Manufacture of the *AMF-AnastomosisChip* master mould and microfluidic device fabrication was conducted as described previously [10]. Briefly, PDMS (10:1 ratio of base to curing agent; Sylgard 184, Dow Corning, USA) was poured onto the master mould after degassing and cured overnight. After removing the PDMS from the mould, it is diced, inlets/outlets are punched and the slabs subsequently bonded to glass-bottomed Petri dishes using a plasma cleaner. Before use, devices were sterilised again under UV light (254 nm) for 30 min. Details of how the AMF spores and fatty acid (FA) salt treatments are loaded into the *AMF-AnastomosisChip* are provided in the Supplementary Methods.

### Image capture and quantification

The inverted microscopes used, and the necessary specifications for large image capture, are as described for the *AMF-SporeChip* [10]. When imaging fluorescein to visualize the capability of the *AMF-AnastomosisChip* to produce diffusion gradients, an LED with a peak wavelength of ca. 470 nm was used. The following filters were employed on the microscope: FITC filter set with Bandpass filter, 465 nm (Omicron-Laserage Laserprodukte GmbH, Germany); Beamsplitter, 495 nm and Emission filter 525/50 nm (500-550 nm; AHF Analysentechnik AG, Germany).

For the experiment, 20 devices (n = 10 MSR- and n = 10 MSR-FA) were imaged daily until 13 days post inoculation (dpi), as well as at 27 and 36 dpi. This produced a single TIFF image of each whole device, which was then analysed in ImageJ/Fiji (version 1.54f; [29]). Loading factors that were recorded were: (i) numbers of lane pockets holding spores (of n = 18 per device: 9 left, 9 right), (ii) locations of the loaded lanes (Top lanes, 1-3, farthest from the outlet; Middle, 4-6; Bottom, 7-9; Fig. 1B), (iii) numbers of spores per respective pocket, and (iv) the orientation of any subtending hypha of spores (*i.e.*, the initial spore orientation). Spores sitting outside of a pocket but in direct contact with spores in the pocket are counted for that pocket; if they are without direct contact but possess a subtending hypha in the respective pocket, they are counted as 0.5 towards that pocket.

Spore traits that were recorded were: (i) the diameter of the largest spore in the pocket, (ii) the number and average diameter of all germinating spores (a diameter for a spore was counted twice if it had two germination sites), and (iii) the numbers of germination sites leading to hyphae that were in lane versus out of lane (*i.e.*, the resulting spore orientation). Hyphal traits that were recorded (for each in-lane hyphae) were: (i) length (including all branches), (ii) numbers of branches, (iii) times of growth stopping and starting, (iv) lane crosses (main hyphae often grew along one wall of the microchannels – a lane cross was observed if they suddenly moved away and physically reached the other wall), (v) reaching the interaction zone and (vi) if they changed direction when entering the interaction zone. Raw hyphal characteristic data was analysed, as well as used to calculate maximum growth rate from 1-6 dpi as well as 7-13 dpi. Hyphal trait data excluding their lengths were also combined to produce an overall exploration score per hyphae (Tab. 1A). Finally, anastomosis scores were produced, which provided an evaluation of hyphal growth near other hyphae that either led or did not lead to anastomosis, producing a non-binary indicator of hyphal intent to anastomose (Tab. 1B).

**Table 1.**
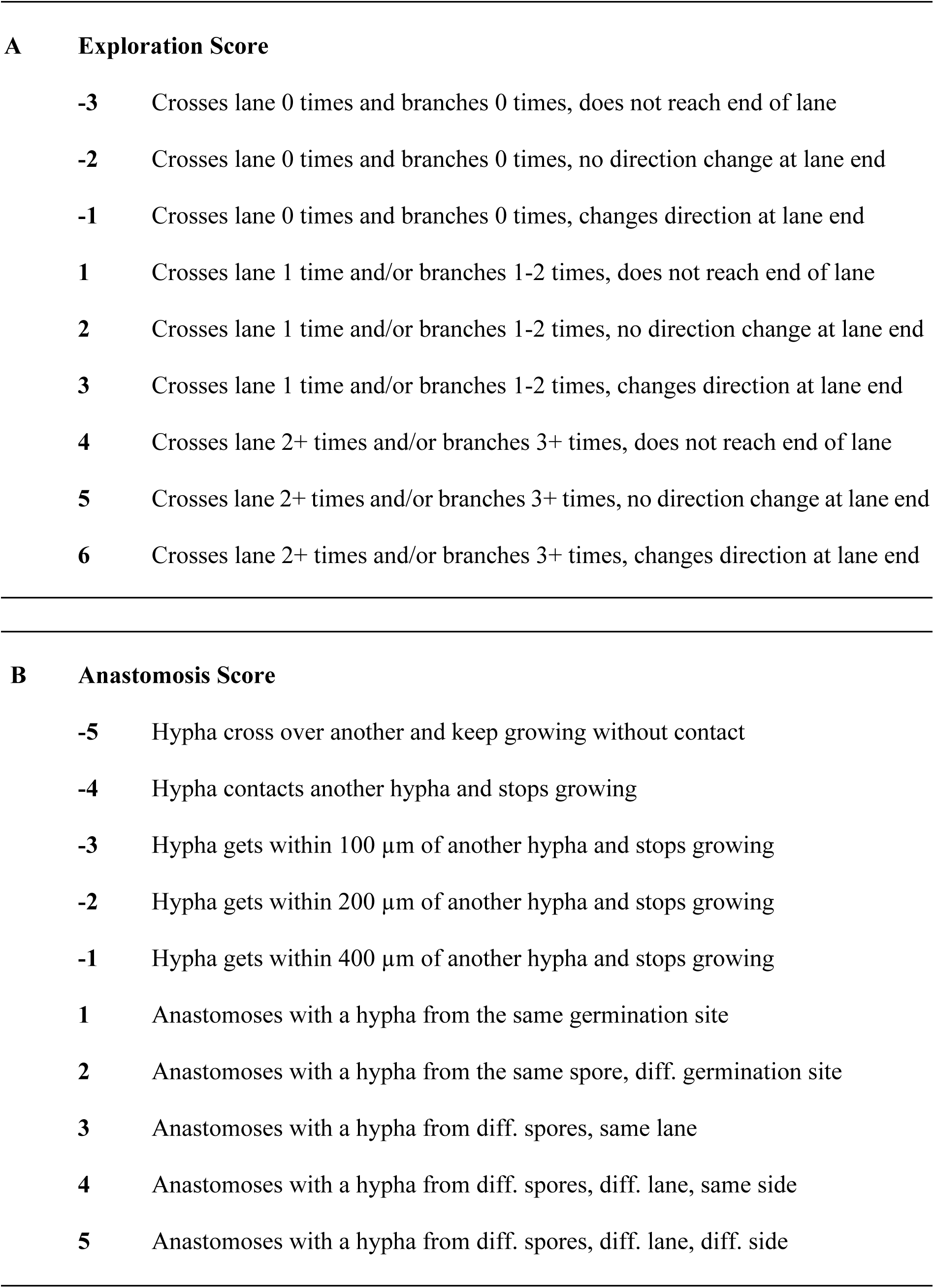
Description of the calculation of the (A) exploration and (B) anastomosis scores. These were produced only for hypha growing from germinated spores. Side refers to the left versus right lanes of the device.

**A Exploration Score**
|  |  |
| --- | --- |
| **-3** | Crosses lane 0 times and branches 0 times, does not reach end of lane |
| **-2** | Crosses lane 0 times and branches 0 times, no direction change at lane end |
| **-1** | Crosses lane 0 times and branches 0 times, changes direction at lane end |
| **1** | Crosses lane 1 time and/or branches 1-2 times, does not reach end of lane |
| **2** | Crosses lane 1 time and/or branches 1-2 times, no direction change at lane end |
| **3** | Crosses lane 1 time and/or branches 1-2 times, changes direction at lane end |
| **4** | Crosses lane 2+ times and/or branches 3+ times, does not reach end of lane |
| **5** | Crosses lane 2+ times and/or branches 3+ times, no direction change at lane end |
| **6** | Crosses lane 2+ times and/or branches 3+ times, changes direction at lane end |
| **B Anastomosis Score** |  |
| **-5** | Hypha cross over another and keep growing without contact |
| **-4** | Hypha contacts another hypha and stops growing |
| **-3** | Hypha gets within 100 $\mu\text{m}$ of another hypha and stops growing |
| **-2** | Hypha gets within 200 $\mu\text{m}$ of another hypha and stops growing |
| **-1** | Hypha gets within 400 $\mu\text{m}$ of another hypha and stops growing |
| **1** | Anastomoses with a hypha from the same germination site |
| **2** | Anastomoses with a hypha from the same spore, diff. germination site |
| **3** | Anastomoses with a hypha from diff. spores, same lane |
| **4** | Anastomoses with a hypha from diff. spores, diff. lane, same side |
| **5** | Anastomoses with a hypha from diff. spores, diff. lane, diff. side |

### COMSOL simulations

COMSOL simulations were used to test the presence of an FA concentration gradient in the *AMF-AnastomosisChip* hyphal interaction zone. FA diffusion was simulated using COMSOL Multiphysics version 6.1, with the transport of dilute species add-on module. The modelled geometry contained 5 mm-high plugs extruding from the inlets and outlets and did not include AMF spores or hyphae. The initial conditions were set to 0.5 mM solute in the outlet plug and 0 mM in the rest of the device (Fig. S1A). We assumed a constant temperature of 27 °C and used COMSOL’s inbuilt “water, liquid” as a global material. We also assumed no-flux external boundaries, so that solute transport occurred only by diffusion. Solute transport within the device was simulated for 13 days for 10 diffusion coefficient (Dc) values ranging from 10-11 to 10-8 m^2^/s, using a relative tolerance value of 0.005. We approximated the Dc of FA in water to be ≈ 4.6×10-10 m2/s by using the experimentally derived Dc of fluorescein, a molecule of comparable size [30]. Solute concentration as a function of distance was derived from concentration values across a line beginning at lower edge of the FA plug and ending at the top of the hyphal interaction zone, with a constant height of 5 µm (Fig. S1B).

Our simulations suggested that an FA concentration gradient was formed in the hyphal interaction zone and persisted for at least 13 days (Fig. S2). The slope of the concentration gradient decreased with time and depended on the solute’s diffusion coefficient (Dc). As may be expected, solutes with lower diffusion coefficients formed steeper concentration gradients (Fig. S1C-D).

### Statistical analysis

The statistical analysis of the obtained data was performed in R version 4.2.0 (2022-04-22) using RStudio (version 2024.09.1+394). Analyses of variance (ANOVAs) were run after a graphical check for homoscedasticity and normality of the trait data when fit to a linear model. Variance decomposition analyses (repeatability) were made with R function *rptR* (package rptR, [31]). These analyses produce the proportions and significances of a trait’s data that are explained by all tested factors as specified in a mixed-effects model. Only non-nested factors are included in the repeatability analyses of a trait.

Repeatability was only performed for particular dpi of trait data, determined as targets from the total germinated spore data: 2, 5, 8, and 12 dpi. More precisely: 2 dpi was chosen because at least some spores had germinated in all devices at this timepoint, including more in control devices. At 5 dpi, spore germination appeared equal among treatments. At 8 and 12 dpi, treated devices surpassed control devices and maintained its lead in total germination and activity until the end of initial tracking. At 27 and 36 dpi, there were quite a few new anastomoses, so these timepoints were included for the variance decomposition of anastomosis score. If any timepoints were not visualized in the repeatability bar graphs, this is because there were not enough non-zero data points to allow the model to converge, leading to the exclusion of this timepoint.

As repeatability analyses do not detail the direction in which a significant factor influences a trait (*e.g.*, for continuous data, if they are positively or negatively correlated), correlations of factors and traits were produced using the R function *ggpairs* (package GGally, [32]) and were represented schematically for each set of repeatability analysis. If correlations were not possible, boxplots were generated (with ANOVAs conducted as described). These are included as Supplementary Information (Fig. S3-S5).

## RESULTS

### Efficiency of the AMF-AnastomosisChip loading

The *AMF-AnastomosisChip* is loosely based on the *AMF-SporeChip*. Its design prioritizes the introduction of two sets of spores (which can be from the same isolate or different isolates or species of AMF) from opposite ends (inlets) of the device, leading to detailed phenotyping of their growth towards and within a single, central interaction plane (Fig. 1A). These devices enable high resolution dynamic imaging from spore introduction and germination to hyphal confrontation and potential anastomosis. For the herein described proof-of-concept experiment, only one AMF strain (*R. irregularis* MUCL 43194) was used and introduced into both inlets of the device.

The strategy to trap the *R. irregularis* spores in pockets suitable for the spore size of this species (20–160 μm) without them entering the lanes nor interaction zone (both at 2–12 μm) was successful. No spores passed into the device further than the pockets (Fig. 1C). Spores did not move among pockets even if media was reloaded through the inlets or outlets. Though not used in this study, some treatments loaded into the outlet could be halted by the height transition from the outlet to interaction zone, leading to special segregation of the treatment from the spores in the devices (Fig. S6).

Loading performance for the *AMF-AnastomosisChip* was analysed in the 20 inoculated devices of the experiment, half without and half with the FA treatment (palmitate and myristate fatty acid salts). There were no significant differences in total numbers of loaded pockets between treatment groups (Fig. 2A). On average, there were 8 pockets loaded with spores per device (44% filled, Fig. 2A, E). Average numbers of spores per pocket were around 2 and did not differ between treatments (Fig. 2B).

**Figure 2.**
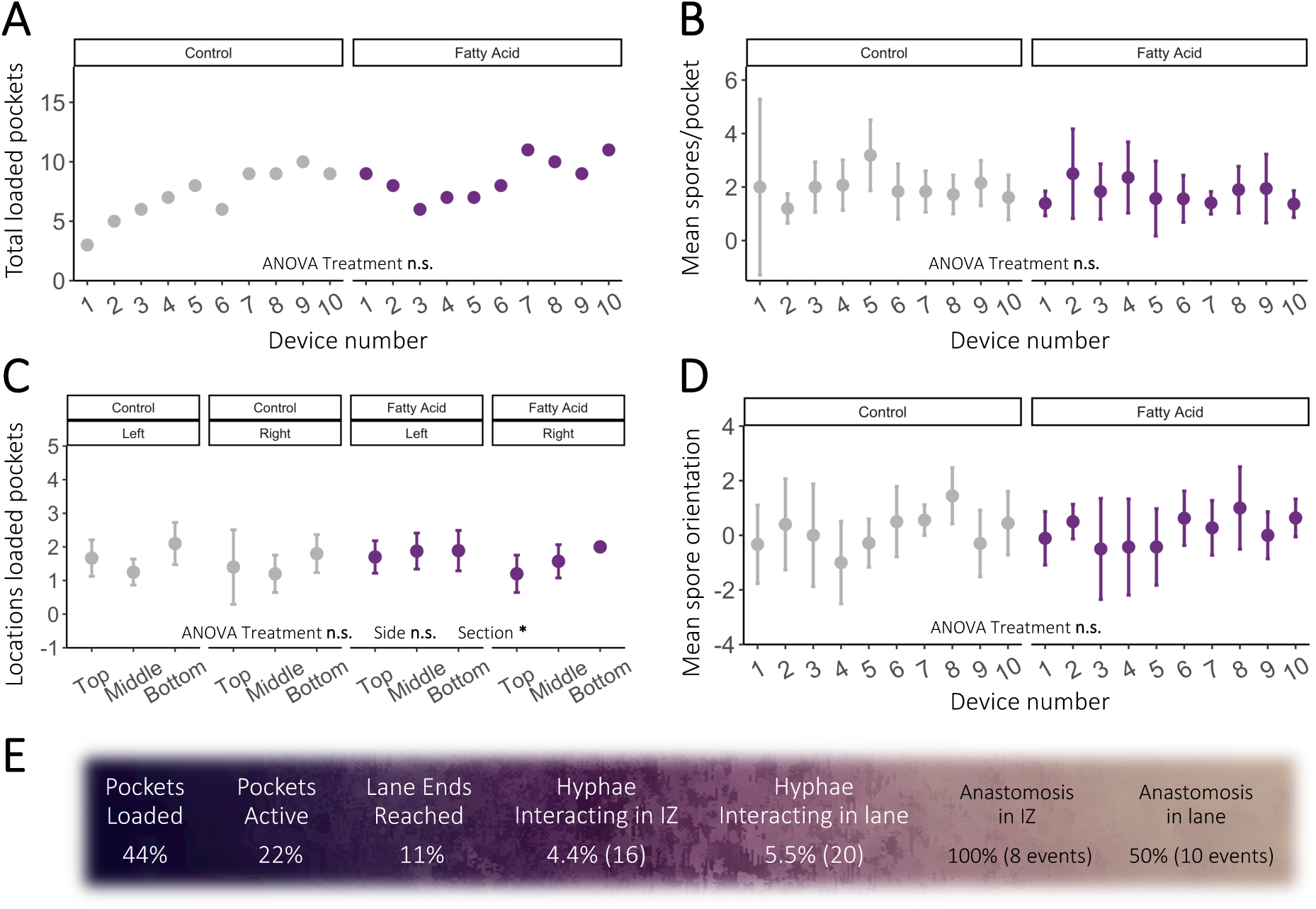
Quantitative analysis of the spore loading performance in the *AMF-AnastomosisChip*. **(A)** Point plot of total loaded pockets per device in each treatment (18 possible pockets; mean_Control_ = 7.2 ± 1.6; mean_FattyAcid_ = 8.6 ± 1.23; n = 10/treatment). **(B)** Point plot of mean spores per pocket in each device (± CI) in each treatment (mean_Control_ = 1.96 ± 0.37; mean_FattyAcid_ = 1.78 ± 0.29). **(C)** Point plot of mean loaded lanes (± CI) in the Top, Middle or Bottom three pockets (x-axis, “Section”) on the left or right of the devices (inner facet, “Side”), per treatment. **(D)** Point plot of mean spore orientations (+ = most growing into lane, - = most growing towards inlet) per pocket in each device (± CI) in each treatment (mean_Control_ = 0.14 ± 0.48; mean_FattyAcid_ = 0.16 ± 0.38). Unless otherwise indicated, the x-axis for (A)-(D) is the Device Number, and the graph is faceted by treatment (“Treatment”: Control, grey; Fatty Acid, purple). ANOVA results are reported at the bottom of each graphic: **n.s.** = not significant, * = *p* < 0.05. **(E)** Visualization of efficiency percentages from loading to possible involvement in anastomoses over the 36-day experiment. IZ = interaction zone.

A small loading bias towards pockets closer to the outlet was observed (Fig. 1, 2C). This did not differ by side of the device (left vs. right), nor by treatment (Fig. 2C). The initial spore orientation (as determined by the direction of subtending hyphae of loaded spores) was not different between devices of the different treatments (Fig. 2D). An average spore orientation value around 0 signified that subtending hyphae of loaded spores were generally half facing the inlets and half facing the lanes.

### Device efficiency in creating potential anastomosis events and overall activity

Approximately four spores per device became active during the experiment (50% of those loaded, Fig. 2E). Of the lanes where activity occurred, half of the hyphae reached the end of their lane, entering the interaction zone (50% of active, 25% of loaded; Fig. 2E). A little less than half of the hyphae in the interaction zone encountered at least one other hypha; this allowed an anastomosis score on that interaction to be produced (40% of hyphae in the interaction zone, coming from 10% of loaded pockets; Fig. 2E). Hyphae did not need to enter the interaction zone to contact and potentially anastomose with each other. Several hyphae from the same or from different spores also anastomosed in-lane. In all, of 400 spores loaded, 36 hyphae came within 400 μm of one other hypha, allowing the attribution of an anastomosis score. 16 of these hyphae encountered each other in the interaction zone (44% of potential anastomoses), and 20 of these within lanes (56% of potential anastomoses; Fig. 2E). On average, 1-2 potential anastomoses occurred per device inoculated with 20 spores (10/side).

Spore germination typically occurred within a week of inoculation, with total germinated spores tapering off after 7 days post inoculation (dpi) in both treatments (Fig. 3A). Mean germinated spores per device was significantly lower in the control devices, and increasingly so over time (ANOVA, significant interaction Treatment:Day, Fig. 3B). There were consistently fewer active growth sites in the control treatment (ANOVA significant by Treatment but no interaction, Fig. 3D, E). Most activity in the devices ended by 12 dpi. Most hyphae making it to the lane ends did so at the peak of device activity around 4-6 dpi (Fig. 3H), but mean numbers of hyphae reaching lane ends per device remained consistent up to 13 dpi and did not differ per treatment (Fig. 3I). Therefore, germination and general activity in the devices may have been affected by the FA treatment but growth up to the lane end seems otherwise not to have been influenced.

**Figure 3.**
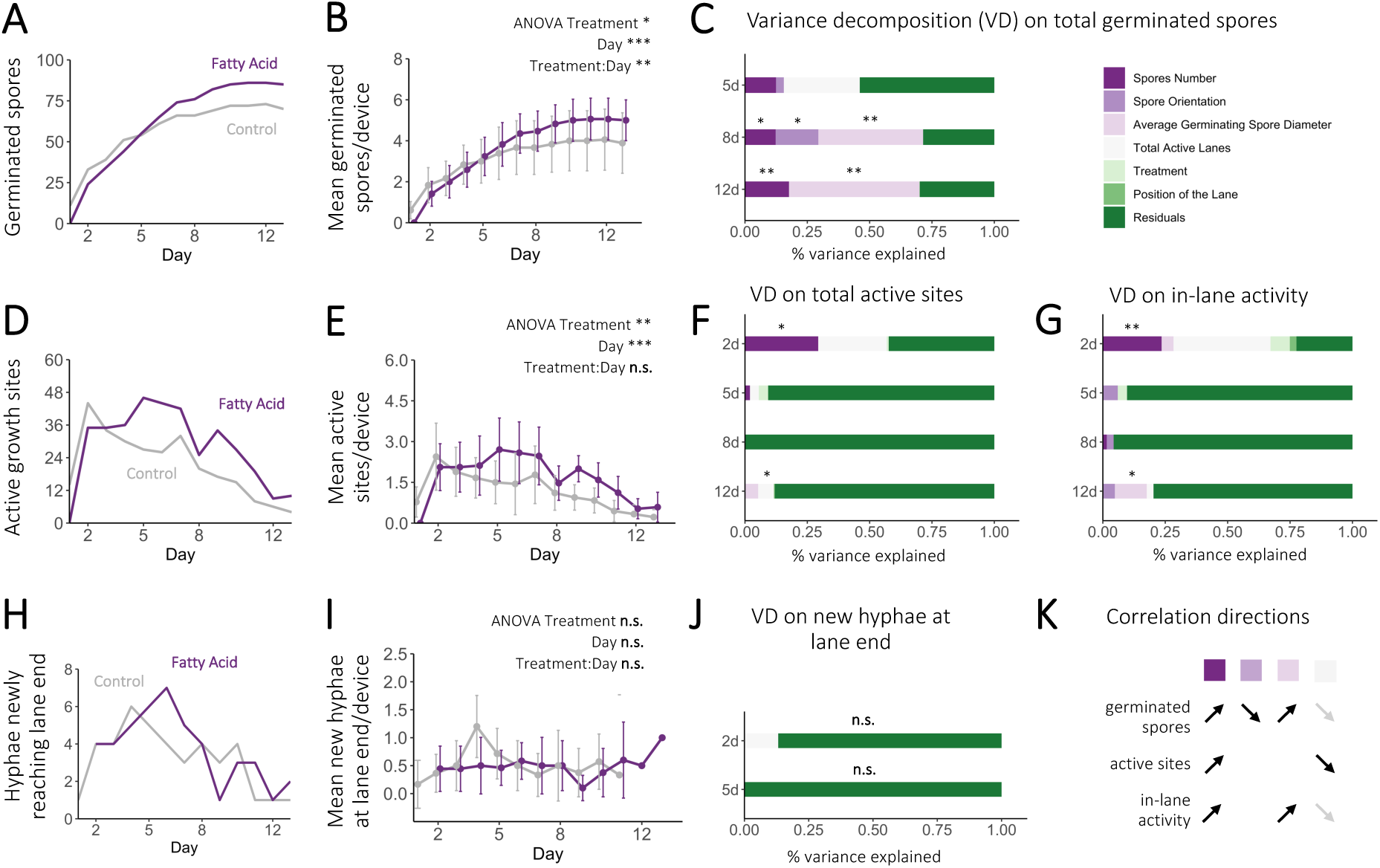
Quantitative analysis of the influences on spore germination and initial growth down lanes. **(A)** Line plot of cumulative germinated spores and **(B)** mean germinated spores per device (± CI) per treatment (Control, grey; Fatty Acid, purple) up to 13 days post inoculation (dpi). **(C)** Variance decomposition (VD) of the cumulative germinated spores in each device, showing percentages of variance explained by seven experimental factors (legend in top right) at 5, 8 and 12 dpi. **(D)** Line plot of cumulative active sites of hyphal growth and **(E)** mean active sites per device (± CI) per treatment up to 13 dpi. **(F)** VD of the cumulative active sites in each device, also including 2 dpi. **(G)** VD of the cumulative active lanes in each device, also including 2 dpi. **(H)** Line plot of total hyphae newly reaching lane ends and **(I)** mean new hyphae at lane ends per device (± CI) per treatment up to 13 dpi. **(J)** VD of the cumulative new hyphae at the end of lanes in each device; there was only sufficient data to run this analysis for the 2 and 5 dpi timepoints. **(K)** Visualization of the general direction (positive, up arrow; negative, down arrow) of the relationship between each variance decomposed trait and variables found to explain significant (black) and large (grey) amounts of variance for that trait. ANOVA results are reported at the top of relevant graphics in (B), (E), (I). ANOVA and VD significances are indicated by: **n.s.** = no significances in the entire bar, * = *p* < 0.05, ** = *p* < 0.01, *** = *p* < 0.001. Unmarked VD sections among other marked sections are also **n.s.**

### Holistic evaluation of influences on initial activity do not point to the FA treatment

Variance decomposition (VD) analyses, like repeatability, are different from analyses of variance (*e.g.,* ANOVAs) because they produce the significance of intra-class correlations. This shows how consistent measurements are among grouping factors including planned and unplanned experimental factors, and variance that does not fall in these grouping factors is attributed to the individuality of each organism, *i.e.,* variance among each spore of an isolate (the “residuals”). This allows for a holistic comparison of the drivers of an isolate’s trait variance. An evaluation of the reliability and consistency of the influencing factor in explaining trait outcomes is innately included in VD analyses, but not in ANOVAs.

The VD analyses determined the influence of number of spores in each pocket, their overall initial spore orientation, and their average germinating spore diameter on germination and hyphal growth traits. At the device level, we also tested the contributions of the current number of active lanes, the treatment with FAs, and the position of the growth lane in relation to the outlet. The number of spores in each pocket influenced the total germinating spores, but only after 8 dpi (Fig. 3C). More spores may have allowed (fairly infrequent) late germinations to occur. The quantity of the spores in the pocket had a stronger positive link with numbers of active sites (of which there can be multiple per spore) and hyphal growth in-lane at 2 dpi than with total germinated spores (Fig. 3F, G).

Total germinated spores per pocket were also influenced by spore orientation and average germinating spore size, but also only starting at 8 dpi (Fig. 3C). Thus, the seemingly “final” numbers of germinated spores were significantly correlated to numbers of spores in the pocket, to more of them having old subtending hyphae facing the inlets, and to a larger average germinating spore diameter (Fig. 3C, K). The VD analyses did not pick up many other known factors influencing initial growth activity after 2 dpi, with only a slight indication of larger germinating spore sizes leading to longer in-lane activity and more active lanes reducing active sites (12 dpi; Fig. 3F, G, K). No tested factor was explanatory for whether hyphae reached the lane ends or not (Fig. 3J), and the FA treatment never significantly explained a portion of variance in any trait when included in VD analyses.

### FAs increase the branching and number of active days of each hypha, but other hyphae present in the same lane have an opposite effect

When analyzed by ANOVAs, the FA treatment was found to increase hyphal branching and the total length of hyphae that were still found to be actively growing, and more so over time (significant interaction Treatment:Day, Fig. 4A, S7). Dividing the initial two-week screen into two periods for analysis – 1-6 dpi where germinations and activity appeared to be increasing and 7-13 dpi where they seemed to taper off – FA treatment did not have a significant effect on maximum hyphal growth rates in either period, nor on their mean last day of activity between 7-13 dpi (Fig. S7). It did have an effect on the last day that hyphae were active within the 1-6 dpi period (Fig. S7). The FA treatment did not alter the number of times a hypha crossed a lane (Fig. 3G), nor each hypha’s exploration score, which resumes the total movement of a hyphae in lane (crosses, branching, length, time of activity; Tab. 1A, Fig. 4G, I).

**Figure 4.**
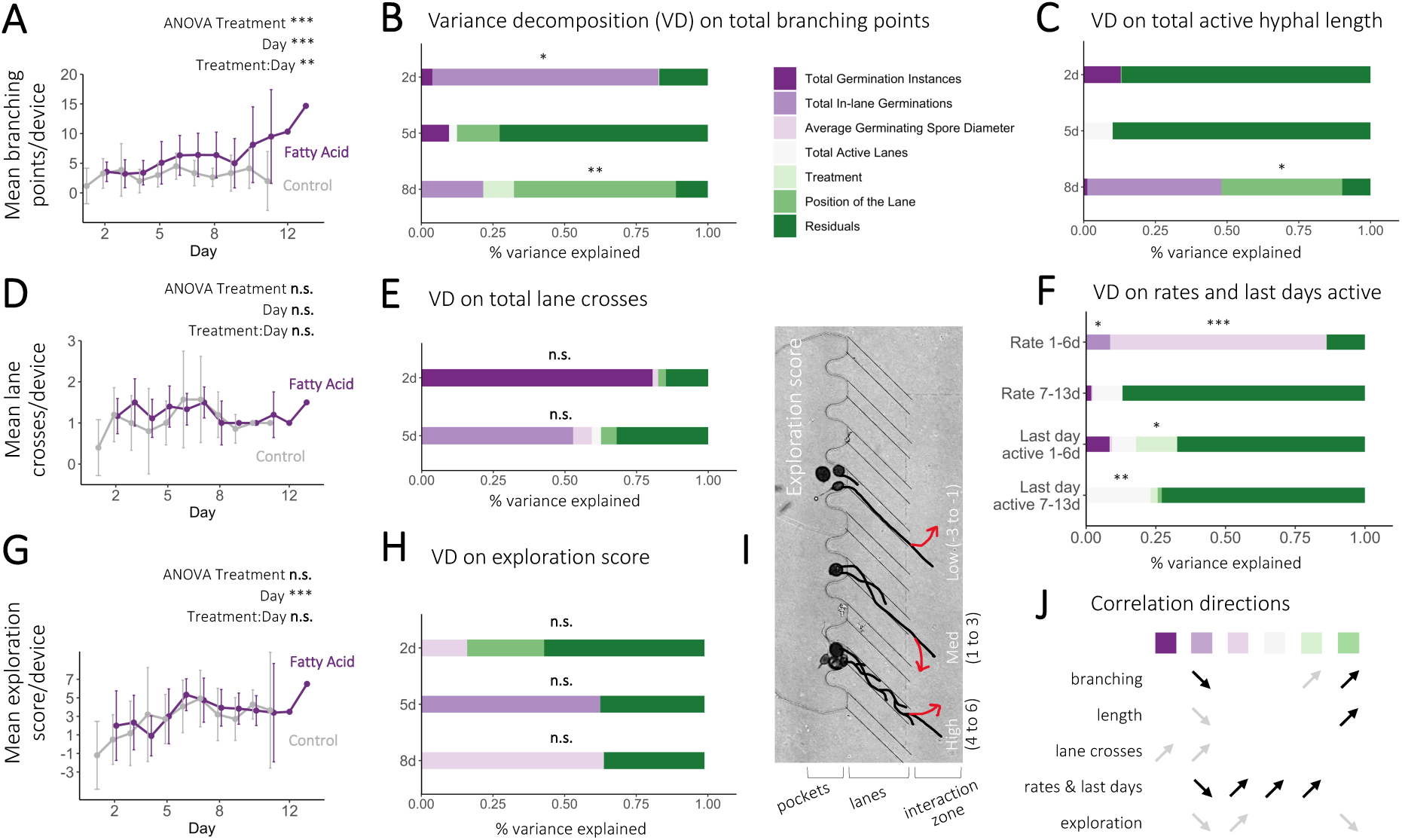
Quantitative analysis of the influences on hyphal lane exploration. **(A)** Mean hyphal branching points per device (± CI) per treatment (Control, grey; Fatty Acid, purple) up to 13 days post inoculation (dpi). **(B)** Variance decomposition (VD) of the cumulative branching points in each device, showing percentages of variance explained by seven experimental factors (legend to the right of B) at 2, 5 and 8 dpi. **(C)** VD of the cumulative length of active hyphae at 2, 5 and 8 dpi. **(D)** Mean hyphal lane crosses per device (± CI) per treatment up to 13 dpi (excludes branching). **(E)** VD of the cumulative lane crosses in each device at the 2 and 5 dpi timepoints, when there was sufficient data to run the VD analysis. **(F)** VD on the maximum growth rate per hypha, as well as the last day that each hypha was active between 1-6 d and 7-13 d. **(G)** Mean exploration score per device (± CI) per treatment up to 13 dpi. **(H)** VD of the cumulative exploration scores in each device at 2, 5, 8 and 12 dpi. **(I)** Schematic to visualize the exploration scores given to each active hypha, which was a combined measure of a hypha’s progression including its branching points, lane crosses, whether it reached the end of the lane, and whether it changed direction at the end of the lane. Low branching/lane crossing scores would be either −3 (did not reach the end of the lane), −2 (reached the end of the lane but did not change direction) and −1 (reached the end of the lane and changed direction, represented by the red arrow). Med (medium) and High branchers/lane crossers were sorted similarly in their respective number ranges (1 to 3 for Med and 4-6 for High). **(J)** Visualization of the general direction (positive, up arrow; negative, down arrow) of the relationship between each variance decomposed trait and variables found to explain significant (black) and large (grey) amounts of variance for that trait. ANOVA results are reported at the top of relevant graphics in (A), (D), (G). ANOVA and VD significances are indicated by: **n.s.** = no significances in the entire bar, * = *p* < 0.05, ** = *p* < 0.01, *** = *p* < 0.001. Unmarked VD sections among other marked sections are also **n.s.**

After running VD analyses, the FA treatment had a positive influence on branching (marginal significance) and on a hypha’s last day of activity between 1-6 dpi (significant; Fig. 4B, F, J). Opposing this, hyphae growing in the same lane (represented by Total In-Lane Germinations in the VD) significantly decreased their early branching and their 1-6d growth rate (Fig. 4B, F, J). Later, these multiple in-lane hyphae decreased active hyphal length, increased lane crosses, and decreased exploration scores (Tab. 1A; Fig. 4C, E, H, J).

At the 2-day timepoint, larger germinated spores also led to higher rates of hyphal growth in the 1-6 d period (Fig. 4F, J). The size of the spore a hypha originated from did not, however, influence the last day the hypha was active; total active lanes in the device, as well as the FA treatment, had more influence in this (Fig. 4F, J). In terms of the positions of these active lanes, a lower lane position (closer to the outlet) increased branching and active hyphal length (Fig. 4B, C, J), following a brief decrease in exploration score at 2 dpi (marginal significance; Fig. 4H, J). The influence of lower lanes on branching could be expected, based on the previously reported role of the FA treatment on hyphal branch occurrences, as well as the previous observation of the FA treatment affecting branching (Fig. 4B). We hypothesized that bottom lanes contained higher FA concentrations than top lanes, as suggested by COMSOL diffusion simulations (Fig. S2).

### Anastomosis was associated with hyphal exploration and occurred more often in the interaction zone

The AMF-*AnastomosisChip* revealed new findings on the influences of anastomosis within this *R. irregularis* isolate. Hyphal interactions leading up to anastomosis could be observed in great detail, in a single plane, over time. This included particular situations when hyphae would initially not interact but would eventually return to anastomose (Fig. 5A, B), or when anastomosis with old subtending hyphae may lead to new hyphal growth on the other end of the old hypha (Fig. 5C). These could have been analyzed separately for their influencing factors, but in this particular work there were not enough instances of anastomosis to do so.

**Figure 5.**
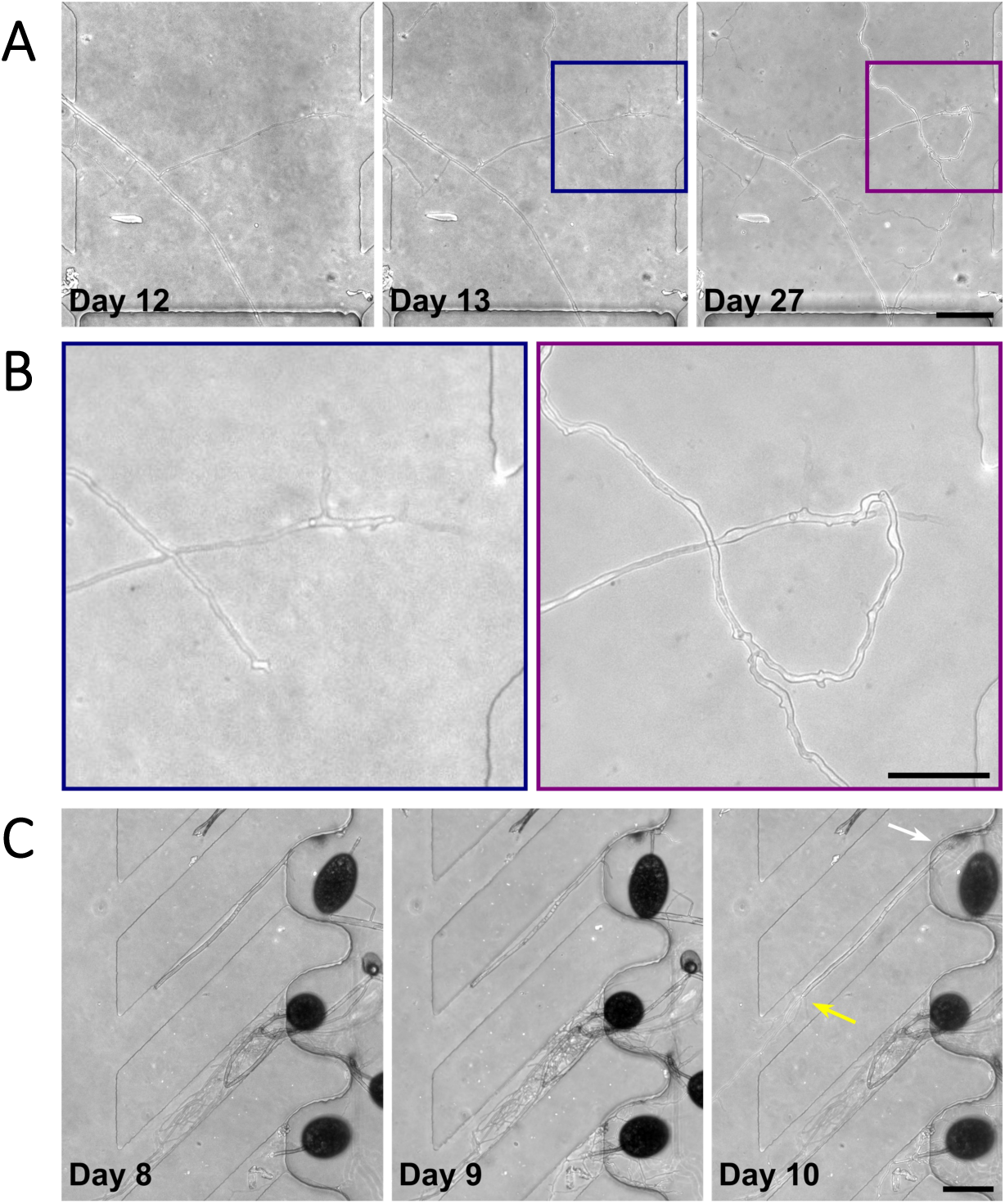
Examples of anastomosis events in the *AMF-AnastomosisChip*. **(A)** Phase contrast time series showing two cross-over events followed by a tip-to-side anastomosis 27 days into the experiment (27 dpi). **(B)** Magnifications of the second cross over from (A), in blue, and the eventual anastomosis, in purple. **(C)** Phase contrast time series showing a spore anastomosing with an old hyphal fragment, using the fragment as a bridge from 8-10 dpi. The white arrow (rightmost panel) indicates the upper end of the hyphal fragment and anastomosis site, and the yellow arrow indicates the lower end of the hyphal fragment, where a fresh hypha emerges following successful anastomosis. Scale bars: 100 µm.

Instances of anastomosis in-lane versus in the interaction zone could be distinguished. Intriguingly, only half of the hyphae that were interacting within a lane ended up perfectly fusing – anastomosing in the form known for intraspecific hyphae (Fig. 2E, 6A). In contrast, every hyphal interaction that occurred among different hyphae, in or near the interaction zone, led to perfect fusion (Fig. 2E, 6A). There was thus a clear difference in interactions among hyphae from the same lane and different lanes.

All events of hyphal interaction were translated to a score on the anastomosis scale which represented whether they especially avoided interaction (*i.e.*, hyphae crossing over each other, the most negative score), stopped growing within shorter to longer distances from each other (somewhat negative to neutral scores), fused with branches of the same hyphae to hyphae in the same lane (neutral to somewhat positive scores), or fused with hyphae from different lanes (the most positive scores; Tab. 1B). If hyphae interacted more than once, an average of their anastomosis scores was produced and used for VD analyses on each hypha. Anastomoses scores were always significantly explained by the exploration score of the hyphae and not at all by FA treatment, position of the lane, and total active lanes (Fig. 6B). This was not just a simple correlation between whether hyphae reached the end of the lane: exploration scores incorporated growth down the lane into each level of exploration (*i.e.*, low scores went from staying in lane to getting to the end, as well as high scores; Tab. 1A).

**Figure 6.**
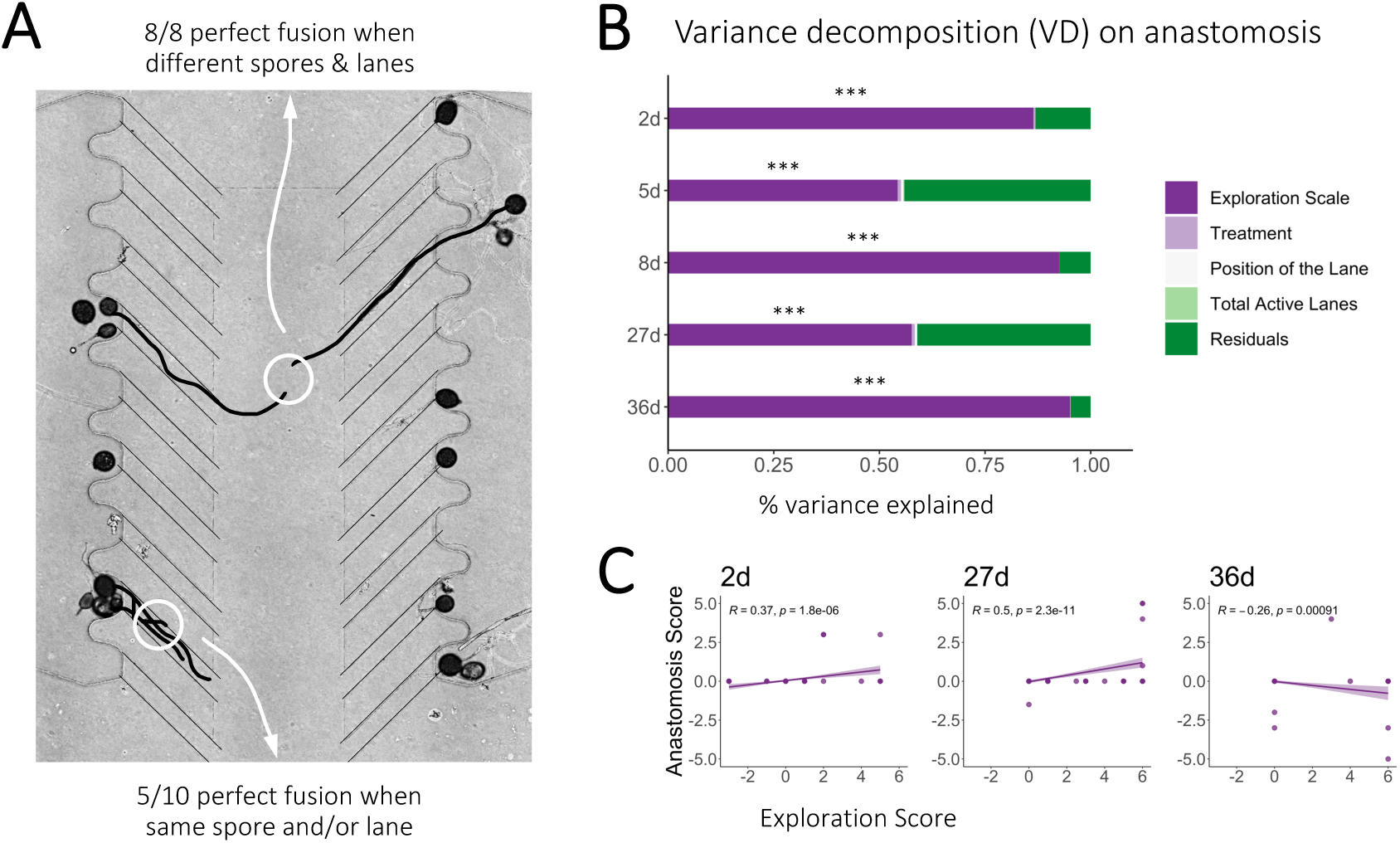
Quantitative analysis of the influences on hyphal confrontations and anastomosis. **(A)** Schematic to visualize different types of anastomoses observed in the *AMF-AnastomosisChip*. All anastomoses were perfect fusions, but only half of hyphal confrontations in-lane resulted in anastomosis (bottom left of the panel, white arrow to numerical summary) while all confrontations in the interaction zone resulted in perfect fusion (top center of the panel, white arrow to numerical summary). **(B)** Visualization of the scale created to score confrontation events for their likelihood of producing an anastomosis event between hyphae originating from different spores. The scale ranges from −5 (where hyphae directly crossed over each other in the shallow lane or interaction space) to 5 (where hyphae perfectly fused when originating from different spores with growth in different lanes). **(C)** Variance decomposition (VD) of the different spore’s anastomosis scale displayed in (B), showing percentages of variance explained by six experimental factors (legend to the right) at 2, 5, 8 and 27 days post inoculation (dpi). At 27 dpi, the devices had just been reloaded with media, but not with the Fatty Acid treatment component. VD significances are indicated by: **n.s.** = not significant, * = *p* < 0.05, ** = *p* < 0.01. Unmarked VD sections are also **n.s.**

From 2 to 27 dpi, exploration scores positively correlated with hyphal interactions ending in perfect fusion (Fig. 6C, left and center panels). Only low and medium explorers produced negative anastomosis scores by 27 dpi. This meant that hyphae that grew fast and did not change directions or branch, even if they ended up in the interaction zone, may have been in a different state of growth, possibly not suitable for anastomosis and thus leading to active avoidance of other hyphae. This study also showed that at 36 dpi, the relationship between anastomosis and exploration was reversed. Hyphae that explored the most then produced the lowest anastomosis scores, demonstrating complete avoidance of anastomosis. Time may be an important component over which the exploration states of hyphae may change.

## DISCUSSION

We used the *AMF-AnastomosisChip* to reveal new findings on asymbiotic AMF spore germination traits (Fig. 1), adding not only a new tool to the field of microbial ecology but also further understanding of how AMF explore and interact. We report that higher local AMF spore densities (*i.e.*, in a single pocket) led to more germination sites, but not necessarily more germinating spores (Fig. 2, 3). In contrast, more active lanes in a device decreased overall growth activity. Larger spores were more likely to germinate and grow faster and longer. Two separate modes of analysis confirmed that a myristate and palmitate fatty acid (FA) treatment increased the time of hyphal activity and numbers of hyphal branches (Fig. 4). The presence of multiple hyphae in a lane generally opposed the FA effect: hyphae grew more slowly, branched less and had lower exploration scores, though they crossed the lanes more. Increased numbers of active lanes conferred effects on hyphal form similar to the FA treatment rather than like in-lane hyphal density. Perfect fusion anastomosis was observed for 50% of in-lane hyphal interactions, but 100% of the time in the interaction zone, among hyphae coming from different lanes (Fig. 5, 6). The higher exploration score a hypha had, the more likely it was to anastomose up to 27 days post inoculation (dpi), after which this was reversed.

Recent literature has called for an increase in high resolution AMF trait phenotyping, both intra- and interspecifically [33,34]. In conjunction, new databases like TraitAM were created to present clean and standardized AMF trait data that can then be reliably analysed to make cross-AMF comparisons [25]. For spore traits, this may be a bit more straightforward, but systematizing the evaluation of asymbiotic traits is challenging due to the obligate nature of AMF and the complications of visualizing hyphal interactions across several planes of solid media. With the *AMF-AnastomosisChip*, high resolution, single-plane, time-resolved observation of spore germination and asymbiotic hyphal activity in liquid media is possible. In addition, hyphae can be screened comprehensively as individuals, in separate lanes, before interacting with other hyphae in-lane or in an interaction zone. This both lays a foundation for future asymbiotic phenotyping while also allowing comparisons to previously published trait information.

Two comparisons to literature were built into the testing of the *AMF-AnastomosisChip*: the number of interactions and anastomoses per spores loaded could be compared to a previous study that characterized monoculture anastomosis events in Petri dishes using the cellophane method [8]. Additionally, half of the devices received an abiotic treatment of FAs (myristate and palmitate) that were previously reported to promote the branching and growth of *R. irregularis* [9]. In Croll et al.’s study, 900 spores were plated per monoculture confrontation, leading to 101-202 hyphal contacts (202-404 hyphae interacting, 22.4%). Approximately half had no interaction, and the other half led to perfect fusion anastomosis (45.8-51.0%). Here, 36 hyphae interacted, originating from 400 inoculated spores (9%). Thus, the hyphal interaction rate was halved in the *AMF-AnastomosisChip*, but this may not be surprising as observations in separate lanes were a specific intention and may have reduced initial contacts. Importantly, if multiple hyphae grew down a lane and interacted there, we found the same percentage of perfect fusion (50%) as in Croll et al.’s work. We additionally replicated the findings of Sugiura et al. in terms of fast growth, more hyphal branching, long growth in terms of time but not in terms of hyphal biomass, in the presence of the FA treatment. This ascertains the validity of the *AMF-AnastomosisChip* in providing AMF trait data that is coherent with previous growth and anastomosis observations.

A novel observation of AMF hyphal interactions presented here is that isolated hyphae that contacted new hyphae from spores in other lanes always formed perfect fusion anastomosis (100%). This was not previously possible to test with other asymbiotic AMF screening methods. Hyphae that interacted in-lane, or in previous set-ups like that from Croll et al., may have been half from different spores, and half from germination sites on the same spore, leading to the observed 50% perfect fusion events. Alternatively, their germinations in the same location may have determined their fusion rates. As we traced hyphae back to shared origin spores, we were able to show that hyphae from different germination sites of the same spore never anastomosed (*i.e.*, no anastomosis scores of 1 were detected, Tab. 1B, Fig. 7). The knowledge that single spore lines produced from single *R. irregularis* isolates can be different genetically [35] and can produce drastically altered plant phenotypes [36,37] may support that perfect fusion in single isolates is prioritized among hyphae from different spores, but not from the same spore. There were some indications of this in previous work [22]. This could be a mechanism to ensure DNA exchange, even within the same isolate, which may be an important insight considering that to date, *R. irregularis* appear asexual [38] and would need different mechanisms for maintaining genetic diversity.

AMF hyphae have been characterized to have different roles depending on their growth forms, although this has largely been characterized in the symbiotic phase. Long hyphae without branches or other interruptions are referred to as runner hyphae. Highly branched hyphae are regarded as absorbative [39]. The formation of branching points to contour obstacles was recently reported in microfluidic devices, as well as the intricacies of cytoplasmic retraction in dead end spaces [10,23]. Chemical treatments have also been shown to mainly increase branching (strigolactones, [40]; FAs, [9]; monoterpene glucosides, [41]). To our knowledge, it has not yet been demonstrated that the presence of nearby hyphae can also fundamentally alter branching and exploration in a focal hypha. Most interestingly, our observations present multiple in-lane hyphae as a negative influence on branching, enabling exploration through more lane crosses and slow progressions through available space instead. Runner-like hyphae were not frequently observed in multi-hyphae lanes, and if they were, they did not anastomose. Hyphae that most covered the spaces they passed through, and not necessarily by branching, had higher likelihoods to anastomose, a finding that also improved our understanding of exploration and hyphal interactions.

Decreased likelihood of AMF hyphae to anastomose over time is not necessarily surprising. Previous studies have indicated both decreased hyphal activity and fusions with more time since germination [10,20]. These could be directly related: hyphae may encounter resource deficiency after more time in a closed system, leading to decreased growth and interaction. The *AMF-AnastomosisChip* allowed us to dissect this to show that the two traits are more likely independent. At 36 dpi, almost all hyphal interactions included crossovers (and thus, maintained hyphal activity while directly avoiding fusion) or hyphae halted relatively close to each other (within ∼100 µm). We add an additional observation: though exploration score (more lane crossovers and coverage, directional changes at the end of the lane) at first correlated with the likelihood of anastomosis, at 36 dpi, the more that an active hypha had explored to that point, the more likely it was to now avoid fusing with other hyphae, even from other lanes. Despite still having the resources to be active, these hyphae may now have been reprioritized for other functions (*e.g.*, faster host localisation) after certain periods of time in an asymbiotic growth environment, also unrelated to FA presence (Fig. 6).

The observations made possible by the *AMF-AnastomosisChip* were dependent on the number of successfully loaded and active lanes. This relies on the quality and quantity of the spore material loaded. In general, loading performance is improved with single spores that have only short subtending hyphae (<100 µm). Unfortunately, this cannot always be guaranteed, as some subtending hyphae are nearly invisible under the stereoscope. Running the spore source solution through a blender prior to selection and transfer into the devices may address this issue. This could help increase the number of lanes loaded in the devices. Less than half of the lanes were currently loaded in the devices, and so the 1-2 anastomoses observed per device could easily be doubled. Increasing loaded spore numbers in the inlets could also help as spore density in the pockets positively affected numbers of germination sites and the probability of in-lane germination and, therefore, activity that could be monitored at high resolution.

Mostly, downsides to the *AMF-AnastomosisChip* were methodological and could be easily addressed. The FA salts used for the MSR-FA devices were virtually insoluble in water, as well as DMSO, ethanol, methanol and acetone. However, applying this treatment within media plugs was a sustainable method that could also theoretically produce single molecules that diffuse into the device to cause responses in the fungus, reminiscent of an experiment described by Baranger and colleagues [42]. More soluble treatments can be easily applied into the devices and can even be more thoroughly modelled for their diffusion times throughout (Fig. S1, S2). Apart from its huge potential for further systemic work on AMF asymbiotic traits, it can be adjusted to different AMF species and intra- and inter-specific confrontations. The appearance, if any, of anastomosis-specific structures in individual and grouped hypha, from the same or different species, can be analysed in detail; the system is compatible with the use of stains that could highlight different AMF structures. It is also efficient in testing the role of abiotic and biotic treatments. This is of great interest for microbial ecologists aiming to precisely explore how AMF hyphal growth and interactions will react to different nutrient environments, plant signals other than FA (*e.g.*, strigolactones and flavonoids), climate-related factors (*e.g.*, increases in CO_2_). Either single components of microbiomes, or mixed communities, could also be introduced into the device inlets or outlets. This could allow researchers to elaborate on asymbiotic AMF inoculation effects in more detail than ever possible before, which is a pressing question in the field.

## SUPPLEMENTARY INFORMATION

**Supplemental Methods.** Details on device production, loading and COMSOL simulations.

**Figure S1.** Raw correlated data for Figure 3K.

**Figure S2.** Raw correlated data for Figure 4J.

**Figure S3.** Raw correlated data for Figure 6B not shown in Figure 6C.

**Figure S4.** Fatty acid (FA) treatment microscopic images when in the devices but not in plugs.

**Figure S5.** Raw data and statistics for hyphal length per device over time, and growth rate and last day of a hypha’s activity within the earlier (1-6 d) and later (7-13 d) growth periods.

**Figure S6.** COMSOL simulations of FA diffusion in the AMF-AnastomosisChip.

**Figure S7.** Time-course of simulated FA diffusion.

## ACKNOWLEDGEMENTS

We thank the Department of Bioengineering at Imperial College London and the University of Lausanne for their support of F.R. and E.M, respectively.

## DATA AVAILABILITY

All relevant data are available from the corresponding author upon request.

## AUTHOR CONTRIBUTIONS

Conceptualization: F.R., E.M., M.C.-S., K.C., S.D., I.S., C.E.S.

Methodology: F.R., E.M., M.C.-S., K.C., S.D., I.S., C.E.S.

Formal Analysis: F.R., E.M.

Investigation: F.R.

Resources: S.D., I.S., C.E.S.

Data Curation: F.R.

Writing - Original Draft: F.R., E.M., C.E.S.

Writing - Review and Editing: F.R., E.M., M.C.-S., K.C., S.D., I.S., C.E.S.

Visualization: F.R., E.M.

Supervision: I.S., C.E.S.

Project Administration: F.R., E.M., C.E.S.

Funding Acquisition: I.S., C.E.S.

## REFERENCES

1. Bonfante P, Genre A. Mechanisms underlying beneficial plant–fungus interactions in mycorrhizal symbiosis. Nat Commun 2010;1:48.

2. van der Heijden MGA, Martin FM, Selosse M et al. Mycorrhizal ecology and evolution: the past, the present, and the future. New Phytologist 2015;205:1406–23.

3. Brundrett MC, Tedersoo L. Evolutionary history of mycorrhizal symbioses and global host plant diversity. New Phytologist 2018;220:1108–15.

4. van der Heijden MGA, Klironomos JN, Ursic M et al. Mycorrhizal fungal diversity determines plant biodiversity, ecosystem variability and productivity. Nature 1998;396:69–72.

5. Rillig MC. Arbuscular mycorrhizae and terrestrial ecosystem processes. Ecology Letters 2004;7:740–54.

6. Kokkoris V, Lekberg Y, Antunes PM et al. Codependency between plant and arbuscular mycorrhizal fungal communities: what is the evidence? New Phytologist 2020;228:828–38.

7. Basu S, Rabara RC, Negi S. AMF: The future prospect for sustainable agriculture. Physiological and Molecular Plant Pathology 2018;102:36–45.

8. Croll D, Giovannetti M, Koch AM et al. Nonself vegetative fusion and genetic exchange in the arbuscular mycorrhizal fungus *Glomus intraradices*. New Phytologist 2009;181:924–37.

9. Sugiura Y, Akiyama R, Tanaka S et al. Myristate can be used as a carbon and energy source for the asymbiotic growth of arbuscular mycorrhizal fungi. Proc Natl Acad Sci USA 2020;117:25779–88.

10. Richter F, Calonne-Salmon M, van der Heijden MGA et al. *AMF-SporeChip* provides new insights into arbuscular mycorrhizal fungal asymbiotic hyphal growth dynamics at the cellular level. Lab Chip 2024;24:1930–46.

11. Mosse B, Hepper C. Vesicular-arbuscular mycorrhizal infections in root organ cultures. Physiological Plant Pathology 1975;5:215–23.

12. Declerck S. Monoxenic culture of the intraradical forms of *Glomus sp*. isolated from a tropical ecosystem: a proposed methodology for germplasm collection. 1998.

13. Fortin JA, Bécard G, Declerck S et al. Arbuscular mycorrhiza on root-organ cultures. Can J Bot 2002;80:1–20.

14. de la Providencia IE, Fernández F, Declerck S. Hyphal healing mechanism in the arbuscular mycorrhizal fungi *Scutellospora reticulata* and *Glomus clarum* differs in response to severe physical stress. FEMS Microbiology Letters 2007;268:120–5.

15. Rosikiewicz P, Bonvin J, Sanders IR. Cost-efficient production of in vitro *Rhizophagus irregularis*. Mycorrhiza 2017;27:477–86.

16. Jin Z, Jiang F, Wang L et al. Arbuscular mycorrhizal fungi and *Streptomyces*: brothers in arms to shape the structure and function of the hyphosphere microbiome in the early stage of interaction. Microbiome 2024;12:83.

17. Serghi EU, Kokkoris V, Cornell C et al. Homo- and dikaryons of the arbuscular mycorrhizal fungus *Rhizophagus irregularis* differ in life history strategy. Front Plant Sci 2021;12:715377.

18. Oyarte Galvez L, Bisot C, Bourrianne P et al. A travelling-wave strategy for plant–fungal trade. Nature 2025;639:172–80.

19. McGale E, Viray J, Gwyther P et al. Dissecting mycorrhizal fungal trait variation, its genetic basis and trade-offs., DOI: 10.1101/2025.10.08.681103.

20. Giovannetti M, Fortuna P, Citernesi AS et al. The occurrence of anastomosis formation and nuclear exchange in intact arbuscular mycorrhizal networks. New Phytologist 2001;151:717–24.

21. Giovannetti M, Ayio L, Sbrana C et al. Factors affecting appressorium development in the vesicular–arbuscular mycorrhizal fungus *Glomus mosseae* (Nicol. & Gerd.) Gerd. & Trappe. New Phytologist 1993;123:115–22.

22. Giovannetti M, Citernesi AS. Time-course of appressorium formation on host plants by arbuscular mycorrhizal fungi. Mycological Research 1993;97:1140–2.

23. Hammer EC, Arellano-Caicedo C, Mafa-Endara PM et al. Hyphal exploration strategies and habitat modification of an arbuscular mycorrhizal fungus in microengineered soil chips. Fungal Ecology 2024.

24. Richter F, Bindschedler S, Calonne-Salmon M et al. Fungi-on-a-Chip: microfluidic platforms for single-cell studies on fungi. FEMS Microbiology Reviews 2022;46:fuac039.

25. Chaudhary VB, Nokes LF, González JB et al. TraitAM, a global spore trait database for arbuscular mycorrhizal fungi. Sci Data 2025;12:588.

26. Keymer A, Gutjahr C. Cross-kingdom lipid transfer in arbuscular mycorrhiza symbiosis and beyond. Current Opinion in Plant Biology 2018;44:137–44.

27. Chabot S, Bécard G, Piché Y. Life cycle of *Glomus Intraradix* in root organ culture. Mycologia 1992;84:315–21.

28. Cranenbrouck S, Voets L, Bivort C et al. Methodologies for *in vitro* cultivation of arbuscular mycorrhizal fungi with root organs. In: Declerck S, Fortin JA, Strullu D-G (eds.). *In* Vitro Culture of Mycorrhizas. Berlin, Heidelberg: Springer Berlin Heidelberg, 2005, 341–75.

29. Schindelin J, Arganda-Carreras I, Frise E et al. Fiji: an open-source platform for biological-image analysis. Nat Methods 2012;9:676–82.

30. Casalini T, Salvalaglio M, Perale G et al. Diffusion and aggregation of sodium fluorescein in aqueous solutions. J Phys Chem B 2011;115:12896–904.

31. Stoffel MA, Nakagawa S, Schielzeth H. rptR: repeatability estimation and variance decomposition by generalized linear mixed-effects models. Goslee S (ed.). Methods Ecol Evol 2017;8:1639–44.

32. Schloerke B, Cook D, Larmarange J, et al. GGally: Extension to *’ggplot2’*. 2025.

33. Chaudhary VB, Holland EP, Charman-Anderson S et al. What are mycorrhizal traits? Trends in Ecology & Evolution 2022;37:573–81.

34. Antunes PM, Stürmer SL, Bever JD et al. Enhancing consistency in arbuscular mycorrhizal trait-based research to improve predictions of function. Mycorrhiza 2025;35:14.

35. Robbins C, Cruz Corella J, Aletti C et al. Generation of unequal nuclear genotype proportions in *Rhizophagus irregularis* progeny causes allelic imbalance in gene transcription. New Phytologist 2021;231:1984–2001.

36. Angelard C, Colard A, Niculita-Hirzel H et al. Segregation in a mycorrhizal fungus alters rice growth and symbiosis-specific gene transcription. Current Biology 2010;20:1216–21.

37. Ceballos I, Mateus ID, Peña R et al. Using variation in arbuscular mycorrhizal fungi to drive the productivity of the food security crop cassava. 2019, DOI: 10.1101/830547.

38. Lee S-J, Risse E, Mateus ID et al. Evolution of unexpected diversity in a putative mating type locus and its correlation with genome variability reveals likely asexuality in the model mycorrhizal fungus *Rhizophagus irregularis*. BMC Genomics 2024;25:888.

39. Bago B, Azcón-Aguilar C, Goulet A et al. Branched absorbing structures (BAS): a feature of the extraradical mycelium of symbiotic arbuscular mycorrhizal fungi. New Phytologist 1998;139:375–88.

40. Akiyama K, Ogasawara S, Ito S et al. Structural requirements of strigolactones for hyphal branching in AM Fungi. Plant and Cell Physiology 2010;51:1104–17.

41. Tominaga T, Ueno K, Saito H et al. Monoterpene glucosides in *Eustoma grandiflorum* roots promote hyphal branching in arbuscular mycorrhizal fungi. Plant Physiology 2023;193:2677–90.

42. Baranger C, Pezron I, Lins L et al. A compartmentalized microsystem helps understanding the uptake of benzo[a]pyrene by fungi during soil bioremediation processes. Science of The Total Environment 2021;784:147151.

