## Supplementary Information for "Intricate asymbiotic hyphal interactions of *Rhizophagus irregularis* revealed with single-plane observations in a new microfluidic device"

The following Supplementary Information is available for this article:

**Supplemental Methods.** Loading of spores and treatments into the devices.

**Figure S1.** COMSOL simulations of FA diffusion in the AMF-AnastomosisChip.

**Figure S2.** Time-course of simulated FA diffusion.

**Figure S3.** Raw correlated data for Figure 3K.

**Figure S4.** Raw correlated data for Figure 4J.

**Figure S5.** Raw correlated data for Figure 6B not shown in Figure 6C.

**Figure S6.** Fatty acid (FA) treatment microscopic images when in the devices but not in plugs.

**Figure S7.** Raw data and statistics for hyphal length per device over time, and growth rate and last day of a hypha's activity within the earlier (1-6 d) and later (7-13 d) growth periods.

### Supplementary Methods

#### *Loading spores and treatments into the devices*

Spores for the experiments were obtained from split-plate cultures that were at least 3 months old. Under a stereoscope (EZ4 D, Leica, Germany) in a sterilized laminar flow hood, hyphae holding many spores were picked from the fungal compartment with sterilized dissection needles and transferred into a new Petri dish containing liquid MSR- medium (without Phytagel, pH 7). Then, using sterile tattoo needles (03 tight liner, pre-soldered, Barber DTS, UK), the mycelium was cut into single spores with subtending hyphae still attached. Approximately 10 spores were gathered in 21 µl of the MSR- media to be transferred into one of the device inlets and were flushed into the device with a syringe attached to tubing as published previously [1].

The *AMF-AnastomosisChip* differed from the *AMF-SporeChip* in that an opposing, second inlet also had to be loaded with spores. *R. irregularis* spores were prepared and pipetted into this inlet as described for the first. Instead of plugging the tubing (attached to the syringe) into the second inlet, it is inserted instead into the outlet (after pipetting out any liquid in the outlet) and the syringe plunger is pulled outward to build up a negative pressure and aspirate the spores pre-loaded in the second inlet into the pockets located in the Spore Trapping Region (Fig. 1A). To do so, the syringe should be filled with some medium (about 1 ml) as air is compressible/expandable and reduces control over the suction from the syringe. Since the spores tend to follow the air-liquid interface in the inlet, often, the entire medium must be sucked through to introduce all spores into the pockets, resulting in air-filled channels. The outlet syringe is then left plugged in and new medium is pipetted into both inlets and then sucked back into the channels to the outlet tubing. This process can be repeated several times if spores get stuck in the inlet or inlet channel and until liquid fills the device. The tubing should be removed slowly and carefully to prevent a spring-like effect causing spores to bounce out of pockets. Additionally, during all loading steps, tilting the device towards the inlets and tapping the surface will reduce bias towards pockets that are closer to the outlet.

For loading the fatty acid (FA) salt treatment, MSR- solid medium was prepared with supplements of 133 mg/L (0.5 mM) potassium myristate and 147 mg/L (0.5 mM) potassium palmitate (MSR-FA; Alpha Laboratories, UK). After autoclaving, the medium was poured into Petri dishes (~5 mm high). Once solidified and dried, plugs with diameters of 3 mm were punched out and inserted into the outlet of the treated devices (medium was pipetted out of the outlet prior to insertion of the plug). Controls received medium plugs without the supplements. Lastly, all inlets and outlets were topped up with MSR- liquid medium, the Petri dish holding the device was sealed with cling film, and the devices were incubated at 27 °C in the dark.

**Figure S1. COMSOL simulations of FA diffusion in the *AMF-AnastomosisChip*. (A)** A schematic of the modeled geometry with the outlet plug (purple) containing an initial solute concentration of 0.5 mM. **(B)** A top view of the geometry includes a red arrow representing the cut line on which simulated concentrations were plotted as a function of distance in C-D. Simulated FA concentration **(C)** as a function of time, showing a concentration gradient was formed and persisted in the hyphal interaction zone, and **(D)** on day 10 as a function of diffusion coefficients (Dc), showing solutes with low Dc formed sharper concentration gradients. Purple, blue and green regions correspond to the plug, inlet channel, and hyphal interaction zone, respectively.

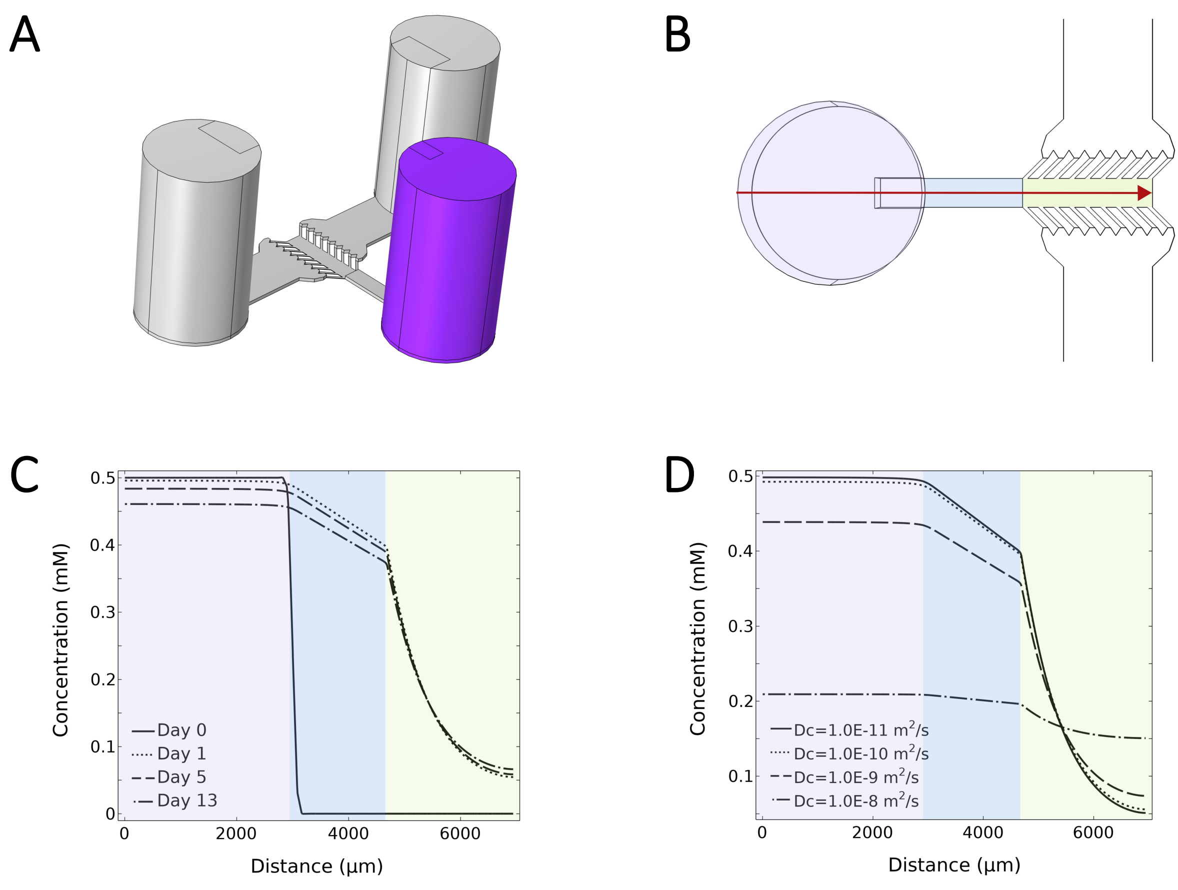

**Figure S2. Time-course of simulated FA diffusion in the *AMF-AnastomosisChip*.** Top view, Dc = $4.6 \times 10^{-10}$ m$^2$/s.

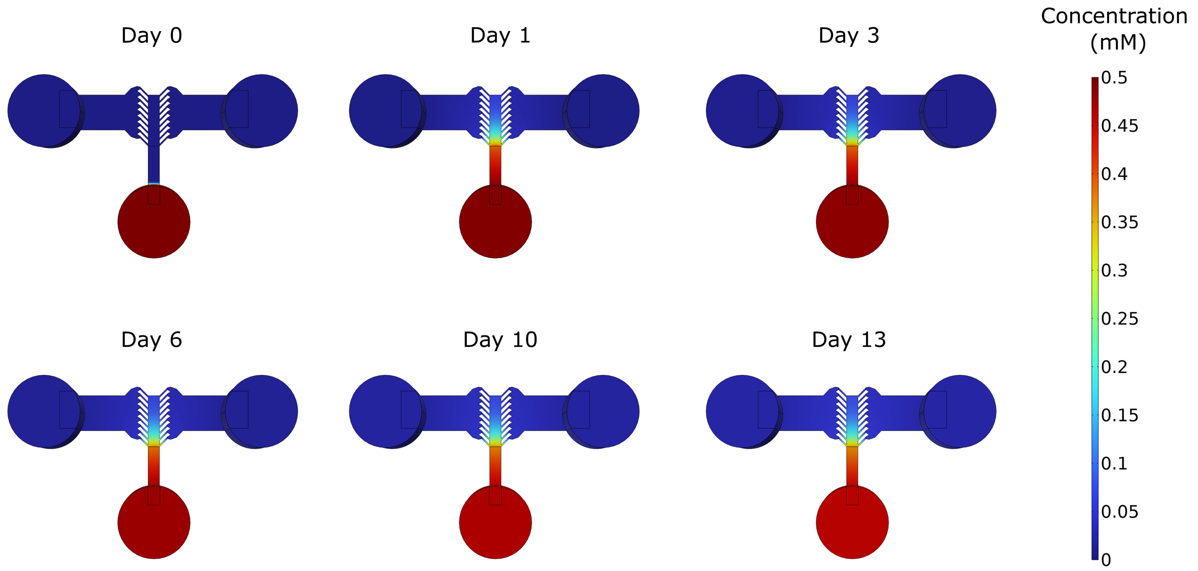

**Figure S3. Raw correlated data for Figure 3K.** Only the direction of influence depicted in Figure 3K is determined by these correlations. Pearson's statistics are left on the correlation panels but may not represent the significance found by VD given the different statistical methods. Panels are arranged by rows corresponding to the measurement time when the significance was detected (*e.g.*, 2, 5, 8 or 12 days post inoculation, dpi). **(A)** Germinated spores (6 panels), **(B)** active sites (3 panels, 2 in-sets representing the trends formed from active lanes on the opposing vs. the same side of the device), **(C)** in-lane activity (3 panels).

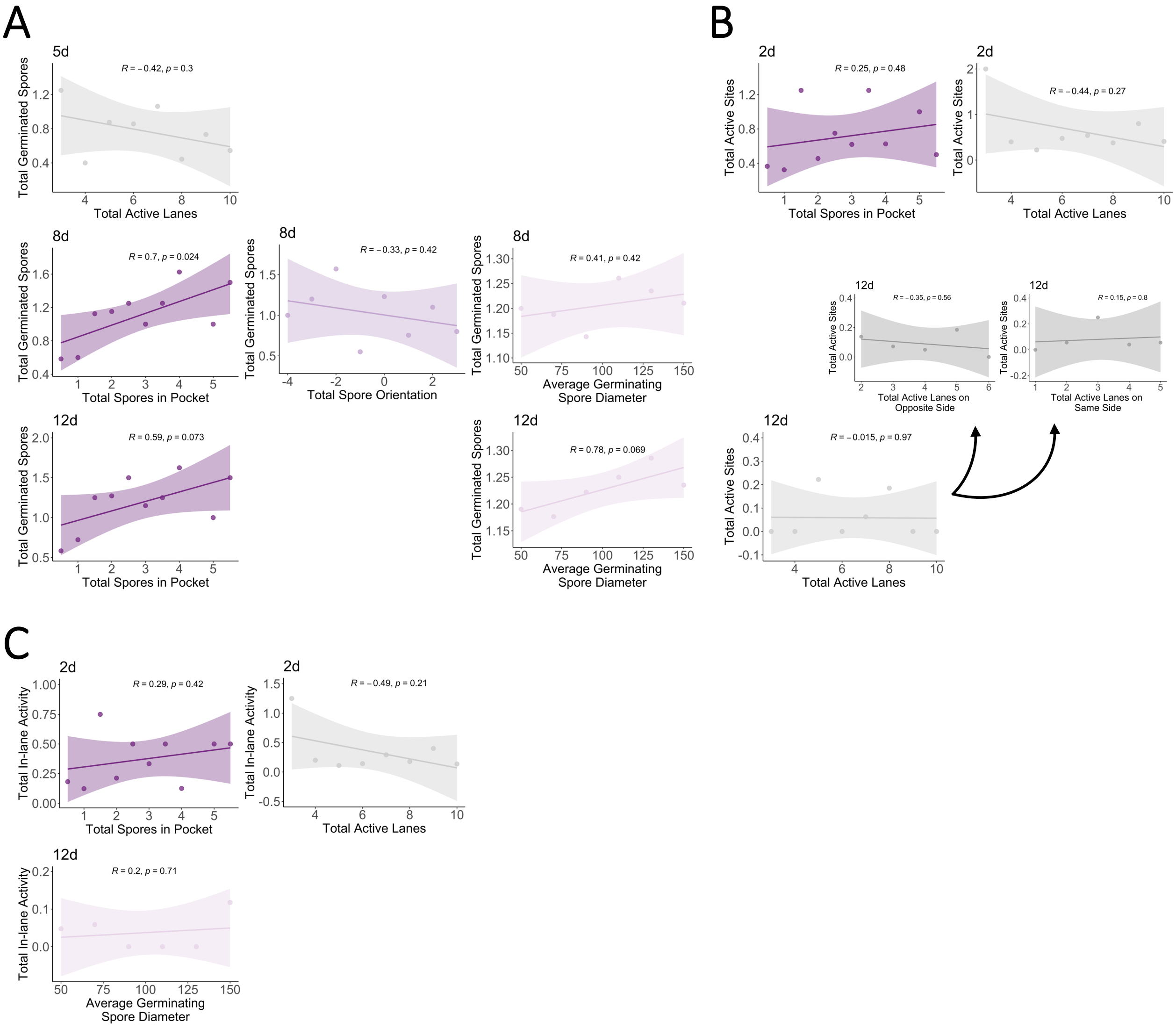

**Figure S4. Raw correlated data for Figure 4J.** Only the direction of influence depicted in Figure 4J is determined by these correlations. Pearson's statistics are left on the correlation panels but may not represent the significance found by VD given the different statistical methods. Panels are arranged by rows corresponding to the measurement time when the significance was detected (*e.g.* 2, 5, 8 or 12 days post inoculation, dpi). **(A)** Total branching points (3 panels), **(B)** total active hyphal length (2 panels), **(C)** hyphal growth rates (2 panels) and last day of activity (2 panels, 2 in-sets representing the trends formed from active lanes on the opposing vs. the same side of the device), **(D)** exploration score (4 panels).

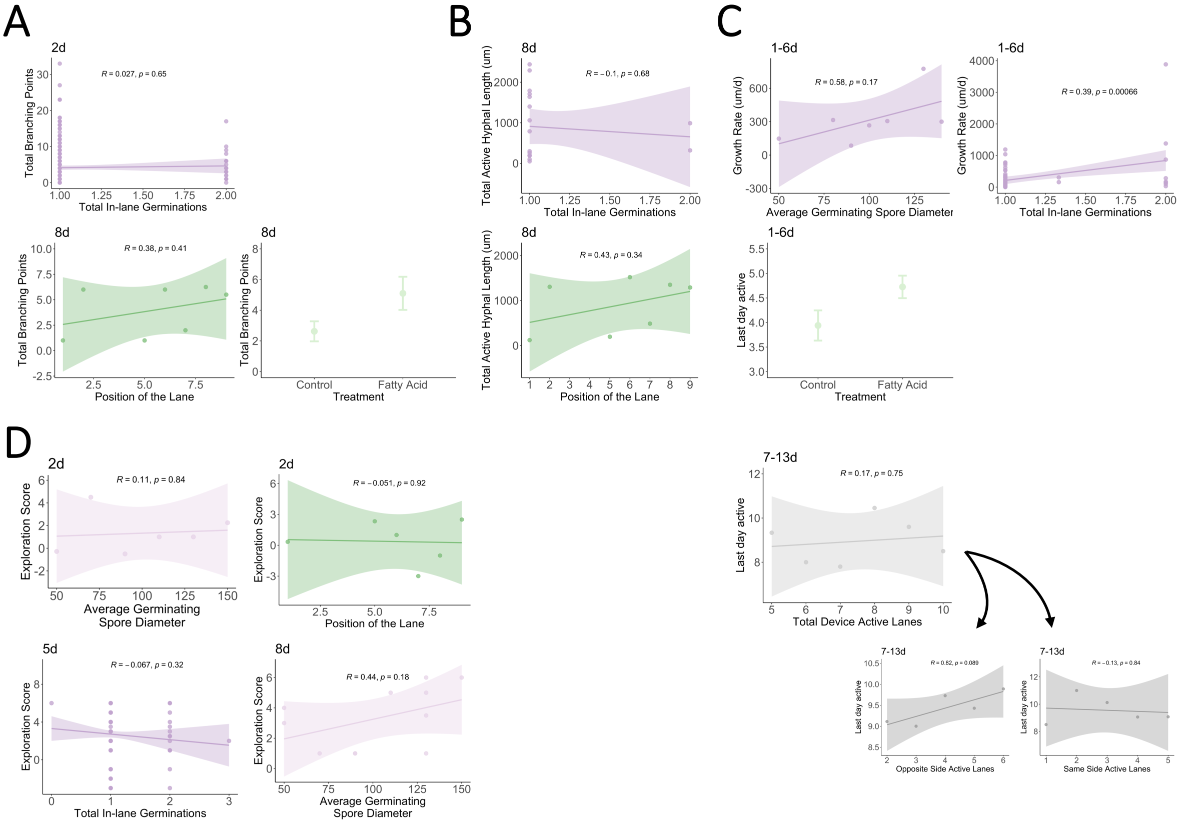

**Figure S5. Raw correlated data for Figure 6B not shown in Figure 6C.** Pearson's statistics are left on the correlation panels but may not represent the significance found by variance composition, given the different statistical methods.

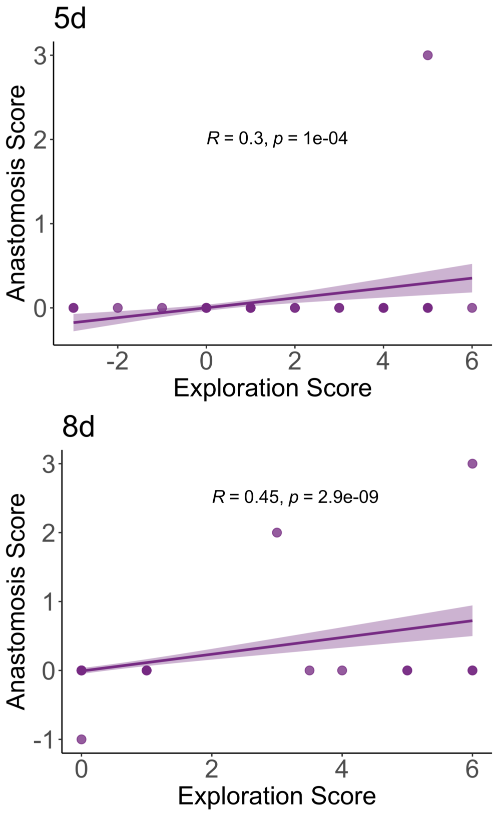

**Figure S6. Fatty acid (FA) treatment microscopic images when in the devices but not in plugs.** **(A)** Mixture of potassium myristate and potassium palmitate, **(B)** only potassium myristate, **(C)** only potassium palmitate. Scale bars: 400 μm.

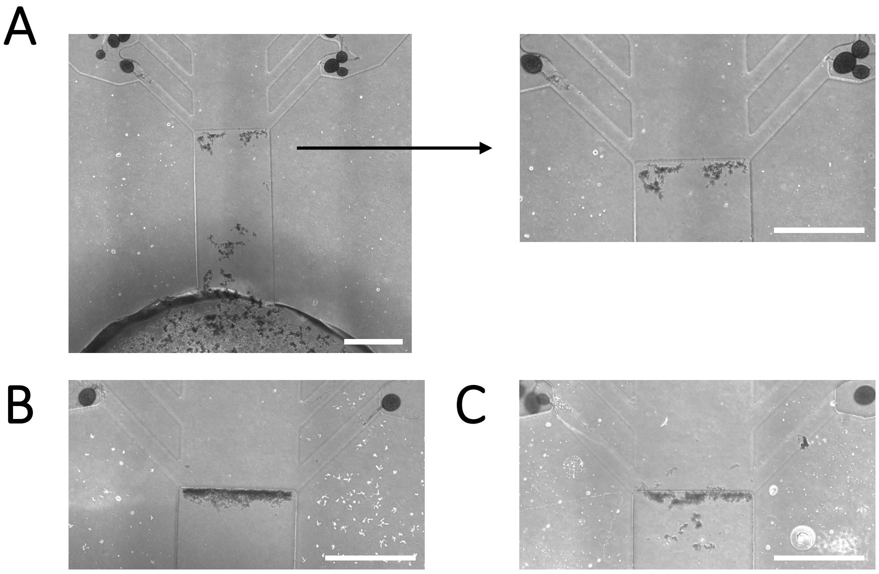

**Figure S7. Raw data and statistics for hyphal length per device over time, and growth rate and last day of a hypha's activity within the earlier (1-6 d) and later (7-13 d) growth periods. (A)** Mean of hyphal length (µm) per device (± CI) per treatment (Control, grey; Fatty Acid, purple) plotted up to 13 days post inoculation (dpi). **(B, C)** Left panels: mean maximum growth rate (µm/d) detected per device (± CI) per treatment during the (B) 1-6 and (C) 7-13 dpi time periods. Right panels: mean last day of activity per hypha, per device (± CI) per treatment during the (B) 1-6 and (C) 7-13 dpi time periods. ANOVA results are reported; significances are indicated by: **n.s.** = no significance, * = $p < 0.05$, ** = $p < 0.01$.

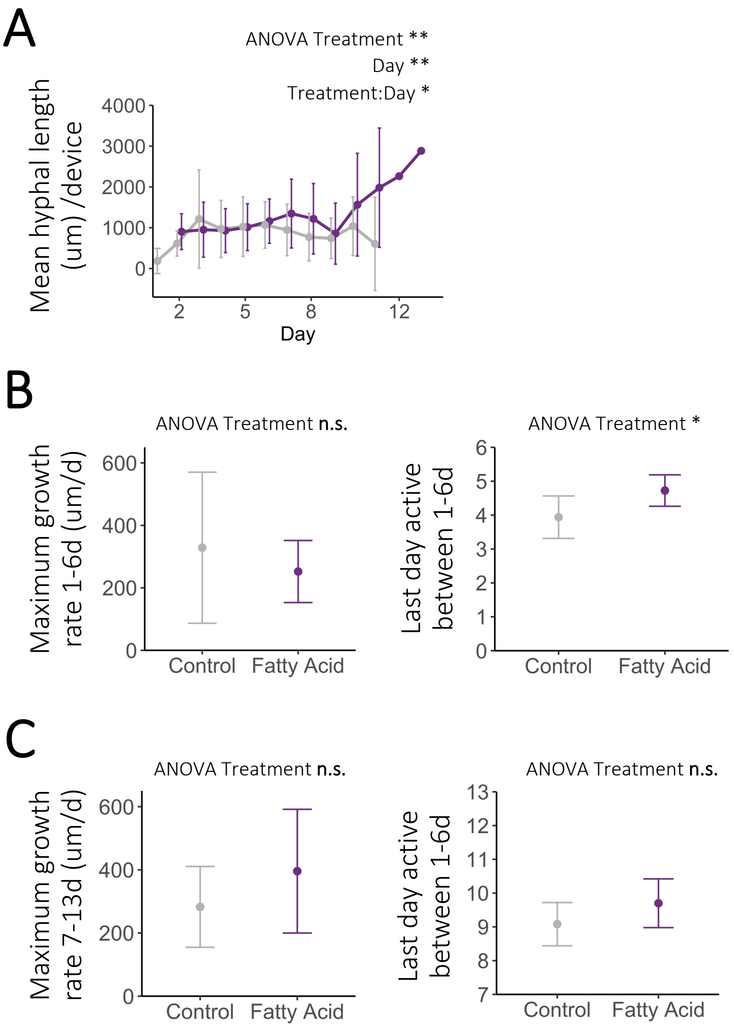

**References**

1. Richter F, Calonne-Salmon M, van der Heijden MGA *et al. AMF-SporeChip* provides new insights into arbuscular mycorrhizal fungal asymbiotic hyphal growth dynamics at the cellular level. *Lab Chip* 2024;**24**:1930–46.
